# Development and molecular characterization of wheat-*Aegilops peregrina* Introgression Lines with resistance to leaf rust and stripe rust

**DOI:** 10.1101/160648

**Authors:** Deepika Narang, Satinder Kaur, Jyoti Saini, Parveen Chhuneja

**Affiliations:** School of Agricultural Biotechnology, Punjab Agricultural University, Ludhiana, Punjab, India

**Keywords:** Aegilops peregrina, Homoeologous pairing, Introgression lines, leaf rust, stripe rust, SSR markers, Wide hybridization

## Abstract

A wild non-progenitor species from wheat tertiary gene pool *Aegilops peregrina* accession pau3519 (UUSS) was used for introgression of leaf rust and stripe rust resistance in bread wheat. It was crossed and backcrossed with hexaploid wheat line Chinese Spring *Ph*^*I*^ to develop two homozygous BC_2_F_6_ wheat-*Ae. peregrina* introgression lines (ILs) viz. IL pau16058 and IL pau16061 through induced homoeologous recombination. Homozygous lines were screened against six *Puccinia triticina* and two *Puccinia striiformis* f. sp. *tritici* pathotypes at the seedling stage and a mixture of prevalent pathotypes of both rust pathogens at the adult plant stage. IL pau16061 showed resistance to leaf rust only while IL pau16058 was resistant to both leaf and stripe rust pathotypes throughout plant life. Molecular characterization of these ILs aided in defining the introgressed regions. Identification of linked markers with advance genomic technologies will aid in marker assisted pyramiding of alien genes in cultivated wheat background.

## Introduction

Wheat is one of the most important staple foods of the world but monoculture cropping of few improved varieties with narrow genetic base made them vulnerable to various biotic and abiotic stresses. Rust diseases significantly reduce wheat production by affecting yield and grain quality (Huerta-Espino et al. 2011, Draz et al. 2015). Wheat leaf rust caused by *Puccinia triticina* Eriks (Pt) develops rapidly at temperature ranging from 10°C and 30°C whereas stripe rust caused by *Puccinia striiformis f. Tritici* Eriks (Pst) appears on wheat grown in cooler climates (2° to 15°C), associated with higher elevations, northern latitudes. Most of the resistance genes have been succumbed up due to the emergence of new races of pathogen through migration, mutation and recombination. Chemical and genetic control are the two methods of rust management but the use of resistant cultivars is the most ideal environmental friendly strategy. Although, the primary and secondary gene pools hold a priority for wheat improvement, additional genes from the tertiary gene pools are anticipated to contribute to sustainable cropping systems (Sears 1981). To date, forty one genes for leaf rust and fifteen for stripe rust resistance have been transferred from wild relatives (Chhuneja et al. 2016, McIntosh et al. 2017). Most of these genes are qualitative and interact with the pathogen in a gene-for-gene manner (Flor 1971). These genes, when pyramided, have the potential to ensure resistance durability across locations. Thus continued effort to transfer genes from *Aegilops* species has been underway to generate genetic materials containing favorable traits in cultivated wheat to be used in future. Furthermore, the development of gene based molecular markers for alien genes will make it suitable for marker assisted transfer of these genes to other elite wheat lines. *Ae. variabils* Eig (syn. *Ae. peregrina* (Hackel in J. Fraser) Maire & Weiller; 2n=4×=28, a tetraploid species with ‘U^p^U^p^S^p^S^p^’ genome (Kimber and Feldman 1987), was thought to be originated by the hybridization between diploid *Ae. longissima* (genome S^l^S^l^) and diploid *Ae. umbellulata* (genome UU) (Kihara 1954, Yu and Jahier 1992). It has been found to be a useful source for resistance to root knot nematode (Yu et al.1990), powdery mildew (Spetsov et al. 1997), stripe rust (Liu et al. 2011) and leaf rust (Marais et al. 2008), making this species an important source to transfer useful genes. Present study describes the transfer of leaf and stripe rust resistance from *Ae. peregrina* accession pau3519 to hexaploid wheat, development of stable introgression lines (ILs), phenotypic screening for rust resistance and their molecular characterization through wheat microsatellite markers.

## Materials and Methods

### Plant materials

A total of thirty one accessions of *Ae. peregrina* (UUSS) are maintained at Punjab Agricultural University, Ludhiana, India. Out of these, twenty three accessions were obtained from National Small Grains Collection (NSGC), Aberdeen, and four were provided by ICARDA, Aleppo, Syria and University of Missouri, Columbia. Leaf and stripe rust inoculums were obtained from Directorate of Wheat Research Regional Research Station, Shimla. These accessions were screened for leaf rust and stripe rust resistance from about last fifteen years. All the accessions were completely resistant to leaf rust while slight variation was observed for stripe rust resistance. A leaf rust and stripe rust resistant *Ae. peregrina* accession pau3519 obtained from USSR, supplied by Dr J. P. Gustafson, University of Missouri, Columbia, USA was found to be resistant to both rusts.

### Development of introgression lines (ILs)

*Ae. peregrina* acc. pau3519, was crossed and backcrossed with hexaploid wheat cv. Chinese Spring (CS) *Ph*^*I*^ stock (Chen et al. 1994) for the homoeologous pairing induction (Fig. 1). The resultant F_1_ plants were crossed with rust susceptible wheat cultivar WL711 (NN), having non necrotic alleles (NN). The F_1_s from this cross were screened against leaf rust and stripe rust pathotypes at the seedling and adult plant stage and the rust resistant plants were backcrossed to WL711 (NN). Leaf and/or stripe rust resistant BC_2_F_1_ plants with plant type similar to WL711 were selfed for five generations and two homozygous wheat-*Ae. peregrina* BC_2_F_6_ introgression lines (ILs) named as IL pau16061 and IL pau16058, having 42 chromosomes were selected at the Field of Punjab Agricultural University (PAU), Ludhiana, India.

**Fig. 1:**
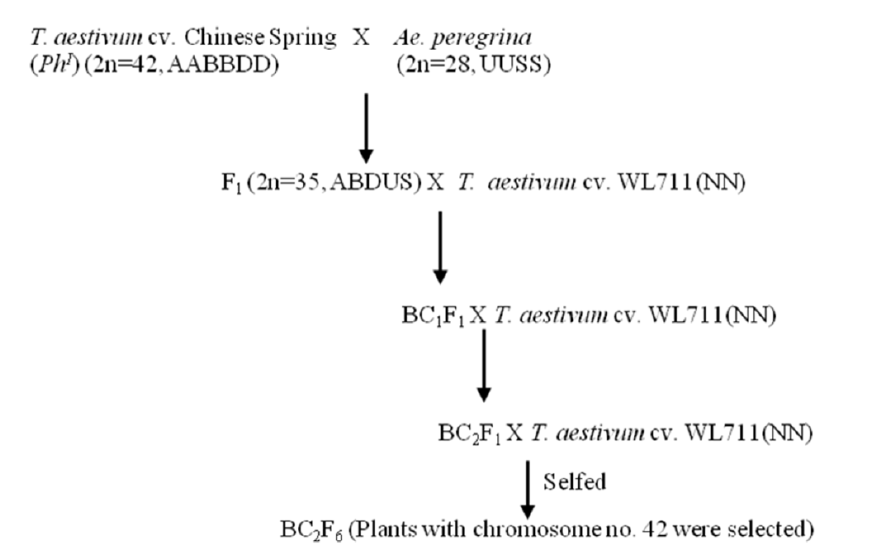
Schematic representation of the crossing strategy adopted for transferring leaf rust and stripe rust resistance genes from *Ae. peregrina* to hexaploid wheat *T. aestivum* cv. WL711 through induced homoeologous pairing using Chinese Spring stock with homoeologous pairing inducer gene, *Ph*^*I*^ gene (*Ph* inhibitor).

### Screening for rust resistance

Rust screening was conducted on BC_2_F_6_ wheat-*Ae. peregrina* ILs alongwith parental lines using six Pt pathotypes viz. 12-5, 77-5, 77-8, 77-10, 104-2 and 104B and two Pst pathotypes viz.46S119 (*Yr9* virulent) and 78S84 (*Yr27* virulent) at the seedling stage. Primary leaves (7-8 days old) were inoculated with urediniospores mixed with talc, kept in humified chambers for 16h in dark and then moved in glasshouse maintained at 18-20°C. Fourteen days after inoculation, infection types (ITs) were recorded on 0-4 scale (McIntosh et al. 1995). The seedlings with infection types 0; to 2 depicts an immune response while 3 or 33+ represents a highly susceptible response. For adult plant assessments, the parental lines were planted in 1.5m rows and spaced 20cm apart. Susceptible checks WL711 and PBW343 were planted around the experimental plot and were sprayed during end December to early January with a mixture of urediniospores of Pt pathotypes (77-2, 77-5 and 104-2) and Pst pathotypes (78S84 and 46S119) suspended in 10L water with few drops of Tween-20. Disease severity was recorded as the percentage of leaf area covered by rust following modified Cobb’s scale (Peterson et al. 1948). Three-four consecutive disease data were obtained to ensure reproducibility of data.

#### DNA extraction and Graphical genotyping

Genomic DNA was extracted from the *Ae. peregrina* acc. pau3519, IL pau16058, IL pau16061, CS(*Ph*^*I*^) and WL711 using the CTAB protocol given by Saghai-Maroof et al. 1994 with minor modifications. DNA was quantified on nanodrop and diluted to a final concentration of 30ng/μl. For molecular characterisation, SSR markers were selected at equal distance from 21 wheat chromosomes from the Composite wheat linkage map at Komugi website (http://www.shigen.nig.ac.jp/wheat/komugi/maps/markerMap.jsp) and the wheat microsatellite consensus map (Somers et al. 2004). PCRs were performed in Applied Biosystems master cycler in reaction volumes of 20μl as described by Roder et al. (1998) with minor modifications. PCR products were resolved either on 2.5% agarose gel or 8% non-denaturing polyacrylamide gel. Molecular marker data were scored as “A” for WL711 specific allele, “B” for *Ae. peregrina* specific allele, “H” for heterozygote and “C” for Chinese spring specific allele. Since introgressions observed were very few, the molecular data of all homoeologous chromosomes (A, B and D) were compiled and represented as group wise. The data was then imported into Graphical Genotype software GGT32 (Berloo 2008) to generate graphical genotype of IL pau16058 and IL pau16061 and to define the size of alien introgressions.

## Results

### Rust Screening

To check the usefulness of rust resistance of wheat-*Ae. peregrina* ILs, six Pt pathotypes viz. 12- 5, 77-5, 77-8, 77-10, 104-2 and 104B and two Pst pathotypes 46S119 and 78S84 predominant in Indian subcontinent, were used to screen this material at both seedling and adult plant stage. At the seedling stage, WL711 and CS (*Ph*^*1*^) exhibited susceptible infection type of 3 or 33+ while *Ae. peregrina* and ILs showed low rust response for leaf rust except IL pau16061 that showed complete susceptibility against both Pst pathotypes.

The same set of lines was tested at adult plant stage against the mixture of leaf rust and stripe rust pathotypes. Cultivar WL711 and CS (*Ph*^*1*^) showed terminal disease severity of 80S and 40S, respectively, whereas ILs and *Ae. peregrina* were highly resistant to leaf rust, indicating leaf rust resistance in both ILs expressed at all stages of plant. Though traces of stripe rust were observed in IL pau16058 but IL pau16061 was susceptible to stripe rust with disease severity of 80S suggesting IL pau16058 carries introgression for both leaf and stripe rust resistance while IL pau16061 encompass seedling leaf rust resistant gene only (Table 1, Fig. 2a &b).

**Table 1.**
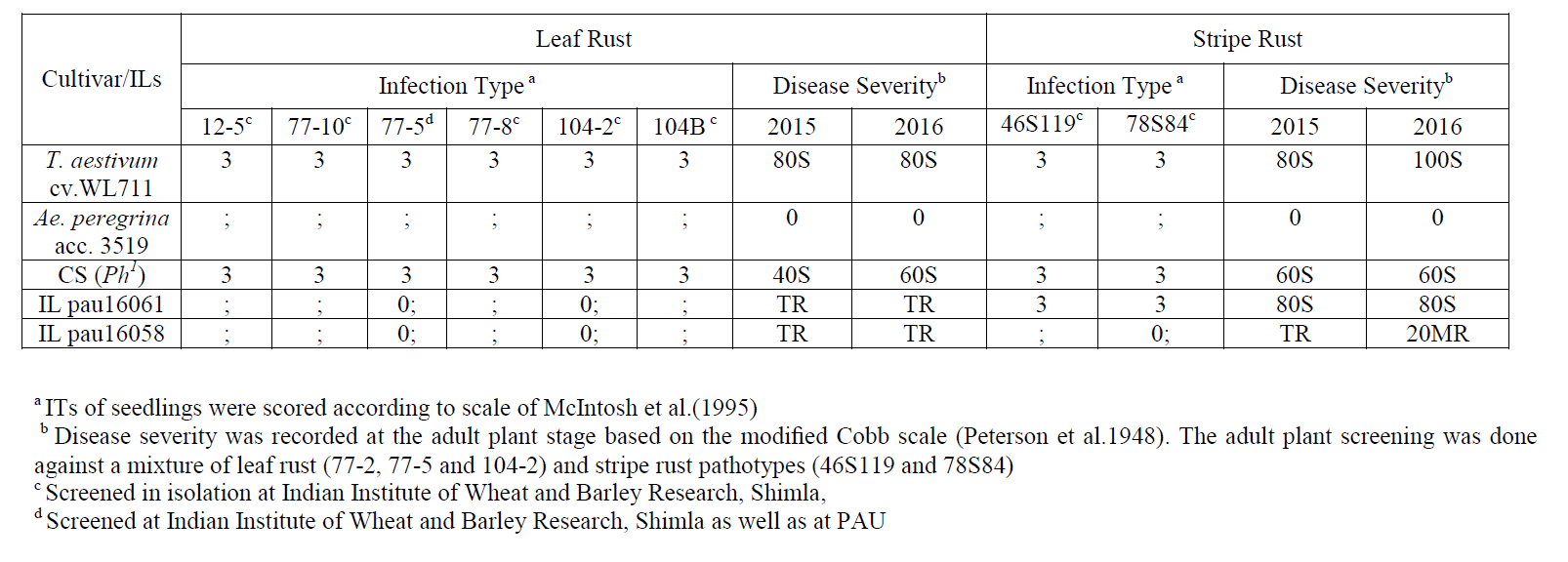
Leaf rust and stripe rust reaction of the selected ILs from the cross *Ae. peregrina* acc. 3519/CS (*Ph*^*I*^) // WL711(NN) at seedling and adult plant stage

**Fig. 2:**
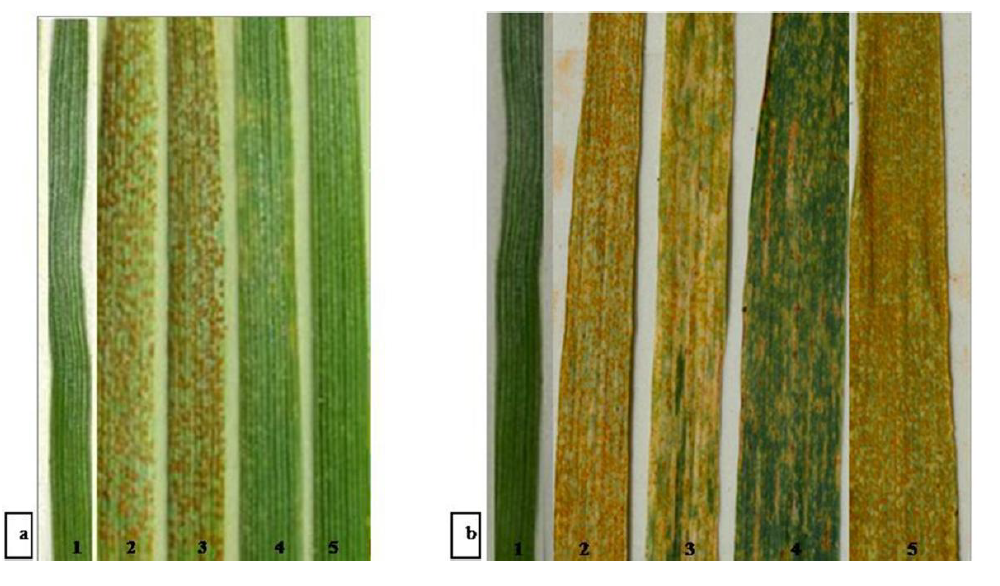
(a) Reaction at the seedling stage against leaf rust pathotype 77–5; (b) Reaction at the adult stage against mixture of stripe rust pathotypes; 1(*Ae. peregrina*); 2 (WL711); 3 (CS(*Ph*^*I*^); 4 (IL pau16058); 5 (IL pau16061).

### **Molecular characterisation of wheat-*Ae. peregrina* ILs**

To characterise *Ae. peregrina* specific introgression (alien segment) in wheat background, genotypes viz. donor wild accession *Ae. peregrina* pau3519, recipient wheat cultivar WL711, ILs and CS (*Ph*^*I*^) were assayed with a total of 350 wheat microsatellite markers selected at regular intervals from all three wheat homoeologous groups. One hundred thirteen markers (32.2%) showed polymorphism between *Ae. peregrina* and WL711. Parental polymorphism was detected on all the three (A, B and D) genomes of wheat. Of 113 polymorphic markers, 26 and 13 markers showed *Ae. peregrina* specific introgression in IL pau16061 and IL pau16058 respectively, that translated into 23% and 11.5% alien introgression in the respective ILs (Table 2). To define the size of the alien introgression, graphical genotype for all the seven wheat linkage groups depicting introgressed regions was generated for both ILs (Fig. 3). In IL pau16061, maximum alien introgression was observed on homoeologous group 6 followed by group 2 while in IL pau16058 maximum alien introgression was observed on homoeologous group 2 followed by group 5. In both ILs, mostly introgressions were localised terminally as depicted on short arm of group 1 and long arm of group 3, 5, 6 and 7 in IL pau16061 and on short arm of group 4, 5 and on long arm of group 1, 3 in IL pau16058. Comparatively, interstitial introgressions were very few as shown on group 7 of IL pau16058 and on group 2 in both ILs. Fourteen SSR markers showed CS specific alleles in both the ILs. The CS specific introgression was detected on all the wheat chromosomes (depicted as light grey areas in graphical genotype).

**Table 2.**
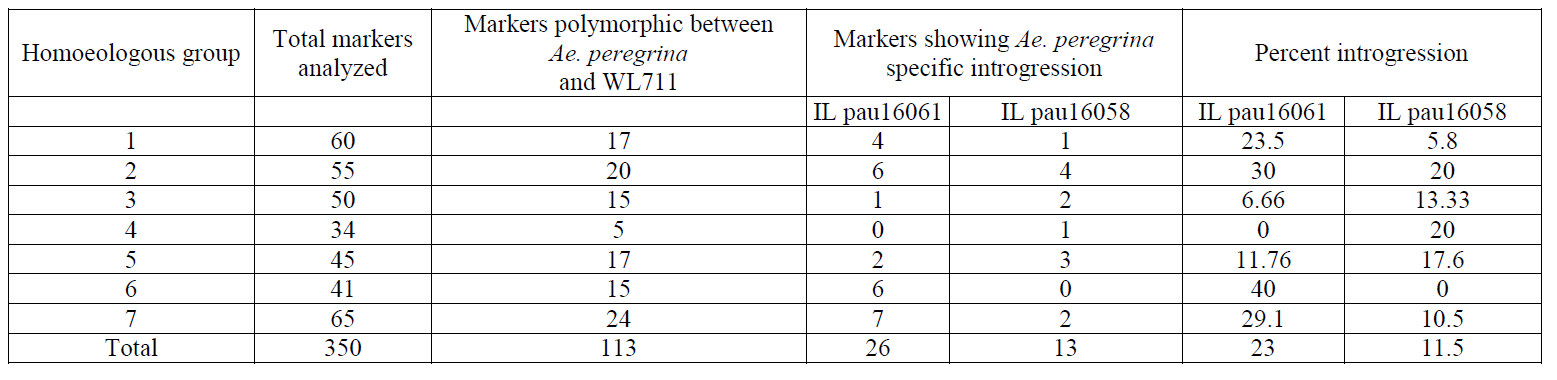
SSR markers showing alien introgression in wheat-*Ae. peregrina* IL pau16061 and IL pau16058

| Homoeologous group | Total markers analyzed | Markers polymorphic between *Ae. peregrina* and WL711 | Markers showing *Ae. peregrina* specific introgression |  | Percent introgression |  |
| --- | --- | --- | --- | --- | --- | --- |
|  |  |  | IL pau16061 | IL pau16058 | IL pau16061 | IL pau16058 |
| 1 | 60 | 17 | 4 | 1 | 23.5 | 5.8 |
| 2 | 55 | 20 | 6 | 4 | 30 | 20 |
| 3 | 50 | 15 | 1 | 2 | 6.66 | 13.33 |
| 4 | 34 | 5 | 0 | 1 | 0 | 20 |
| 5 | 45 | 17 | 2 | 3 | 11.76 | 17.6 |
| 6 | 41 | 15 | 6 | 0 | 40 | 0 |
| 7 | 65 | 24 | 7 | 2 | 29.1 | 10.5 |
| Total | 350 | 113 | 26 | 13 | 23 | 11.5 |

**Fig. 3.**
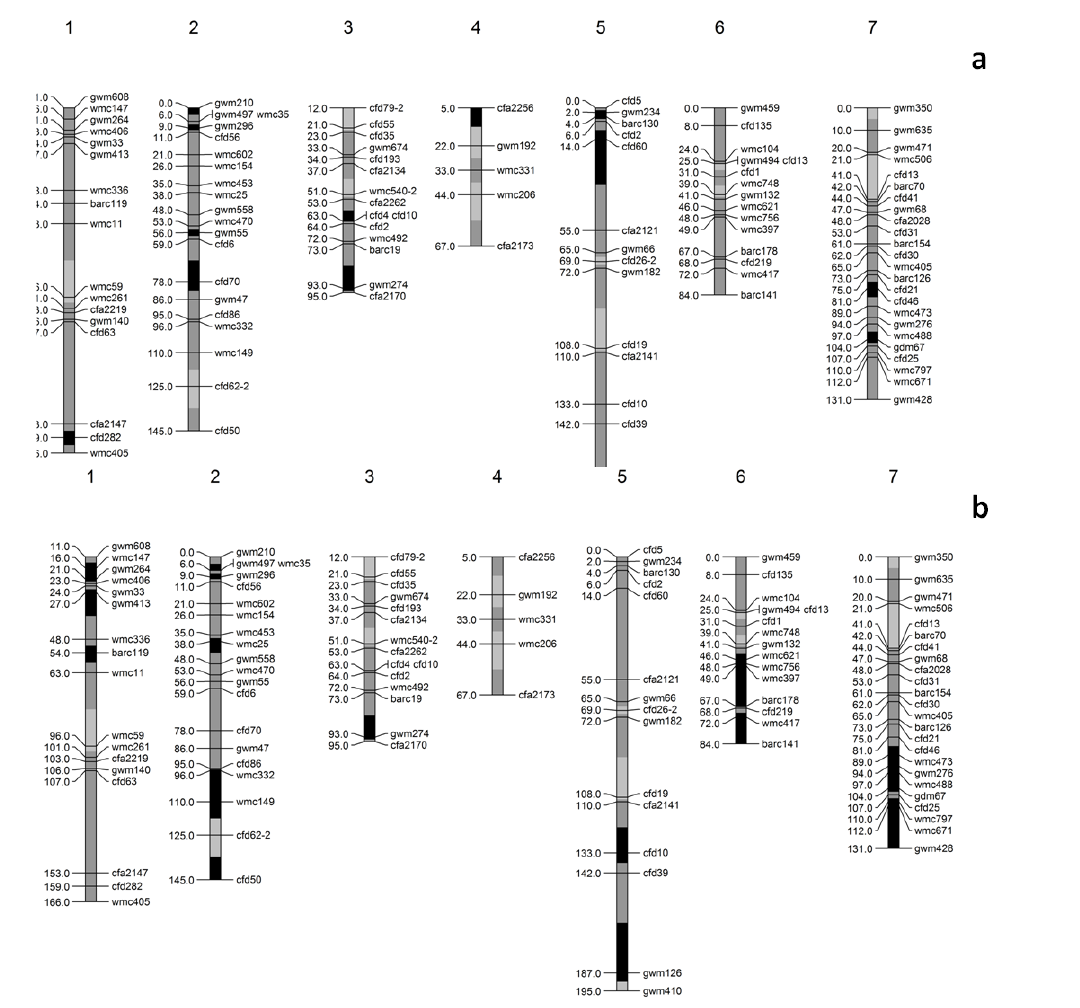
Graphical genotyping of wheat–*Ae. peregrina* introgression lines a) pau16058; b) 16061 using whole genome SSR markers. Grey areas represent wheat-specific alleles and black areas indicate introgression of *Ae. peregrina*-specific alleles. Light grey areas indicate Chinese Spring specific segments. Map distances are according to Komugi composite wheat map

## Discussion

Wheat-alien introgression is the most preferable method to transfer useful traits for biotic and abiotic stress resistance and is a valuable source for considerable genetic diversity to improve yield potential of wheat (Mujeeb-Kazi et al. 2013). Since 1950s, efforts have been made to transfer *Aegilops* chromatin into wheat using ionizing radiation (Sears 1956) or manipulation of *Ph* genetic control system which allows intergenomic pairing (Riley et al. 1968, Chen et al. 1994, Aghaee et al. 2002). In the present study, two wheat-*Aegilops peregrina* ILs have been developed using induced homoeologous recombination. These lines showed resistance to leaf rust and stripe rust due to wheat-alien transfers and chromosome rearrangements occurred during the transfer and selection process. Besides carrying the rust resistant genes, these ILs also harbour new alleles for HMW-glutenin subunits to improve wheat quality (Kaur et al. 2014). None of the IL showed any deleterious phenotypic effects thereby appeared to carry small alien segments with minimum linkage drag indicating introgressions are compensatory.

In Punjab Agricultural University, various addition, substitution, translocation and introgression lines containing alien segments from *Ae. umbellulata*, *Ae. triuncialis, Ae. geniculata, Ae. caudata* have been developed in wheat background (Chhuneja et al. 2016). Some of these translocations comprised entire chromosome arms while few are cryptic introgressions that cannot be detected using cytogenetic–based approaches like *Lr57* and *Yr40* from *Ae. geniculata* and *Lr58* from *Ae. triuncialis* where DNA markers gave an evidence of its introgression (Kuraparthy et al. 2007a,b). Molecular marker technology offers a wide range of novel approaches to characterise minute introgressions. SSR markers are preferably employed in wheat-alien introgression breeding due to its high genome-specificity, whole genome uniform distribution and higher level of polymorphism (Autrique et al. 1995; Brown et al. 1996). The molecular profiling of the wheat-*Ae. peregrina* ILs with the wheat whole genome SSR markers pointed that alien introgressions were mainly terminal and very few are interstitial. The results are in accordance with previous studies stating wheat-alien homoeologous recombination predominantly occurs terminally in gene-rich regions as a consequence of single cross-over events to transfer agronomically useful traits (Qi et al. 2004, Lukaszewski et al. 2005). Besides this, the introgressions were mainly observed on chromosomes B and D of wheat genome. Since ‘U’ genome of *Ae. peregrina* was derived from *Ae. umbellulata* (Zhang and Dvora´k 1992), the probability to transfer alien chromatin from U genome to D genome of wheat is quite high (Zhang et al. 1998). While ‘B’ genome of *T. durum* and *T. aestivum* was originated from ‘S’ genome of diverged member of *Sitopsis* section, hence frequency of transfers from S genome to B genome are more likely. The results indicated the homology of S and U genomes of *Ae. peregrina* with that of B and D genome of wheat. Though SSR markers aided in characterization of regions harbouring alien segments but low marker density impeded the development of high resolution graphical genotypes. To characterise the genes governing the rust resistance, mapping populations have been developed from these ILs. Identification of linked markers with advance genomic technologies will aid in marker assisted mobilization of novel alien genes and hence broaden the narrow genetic base of cultivated wheat germplasm for rust resistance.

Wheat-*Ae. peregrina* ILs also possess agronomically important introgressions that are uncharacterized and will serve as impactful breeding material to transfer novel traits in cultivated wheat.

## Conclusion

The present study described the development of two compensating wheat-*Ae. peregrina* ILs from *Aegilops peregrina* acc. pau3519 using Chinese Spring *Ph*^*I*^ stock through induction of homoeologous pairing. Phenotypic characterization with predominant leaf and stripe rust pathotypes showed IL pau16058 harbours seedling leaf and stripe rust resistance whereas IL pau16061 carries seedling leaf rust resistance only. Molecular profiling with SSR markers defined the regions of alien introgressions to be focussed for characterisation of these novel alien genes that can be used to enhance the narrow genetic base of wheat for rust resistance.

## Acknowledgment

Financial assistance provided by the Department of Biotechnology, Ministry of Science and Technology, Government of India, New Delhi. The provision of rust cultures by Dr. Subhash Bharadwaj, the Indian Institute of Wheat and Barley Research, Shimla is thankfully acknowledged.

